# Capture Efficiency Of Long-Adapter Single-Strand Oligonucleotide Probe Libraries

**DOI:** 10.1101/2023.06.02.543477

**Authors:** Lamia Chkaiban, Lorenzo Tosi, Biju Parekkadan

**Affiliations:** Biomedical Engineering Dept- Rutgers University

**Keywords:** long DNA target cloning, multiplex, Long Adapter Single-Stranded Oligonucleotide, DNA libraries, DNA target cloning, probe target capture optimization

## Abstract

High throughput techniques that can massively produce in parallel, longer DNA sequences of interest can accelerate the decoding of gene functions. LASSO probes are a molecular biology tool that can enrich for DNA targets in a genomic sample via a multiplexed, single-pot reaction for downstream sequencing and/or cloning. Here we have explored aspects of process development and the design of the probes that relate to binding thermodynamics to determine impact on cloned library sequences. Control of ligase concentration, polymerase type, and melting temperature of probe are critical when translating the use of LASSO probes for homogeneous and high fidelity DNA capture.

## Introduction

With the advent of third generation sequencing machines, the cost and time of DNA analysis is going down and the information content continues to expand [1,2]. The increased amount of sequenced organisms has created a complimentary challenge in understanding DNA sequences via functional genomics. Functional genomics attempts to assign a function to predicted genes or regulatory elements to reveal new insights into an organism’s biology [3]. In a more nuanced manner, targeted enrichment in large multiplex formats can also aid in precisely analyzing and expressing structural variants and mutations. The ability to make cloned genetic DNA libraries is of continued importance for DNA sequencing and massively parallel screening assays for functional genomics.

Functional characterization of complex libraries of genetic elements have benefited from flexible, high throughput techniques. Several approaches have been historically developed as a method for selectively cloning a pool of gene in a single experiment. Long range-PCR can be used to encompass a full ORF, though, it is laborious and expensive when dealing with massive libraries of targets a time. To date, DNA sequences of interest are obtained by DNA chemical synthesis as large pool of short DNA sequences. This technique relies on DNA chemical synthesis from releasable DNA microarrays that produce pools of DNA sequences limited in length (maximum ∼300 nucleotide) rarely covering the full length of a gene [4].

<u>L</u>ong-<u>A</u>dapter <u>S</u>ingle-<u>S</u>trand <u>O</u>ligonucleotide (LASSO) probes have been previously developed and tested to capture and clone in parallel over 3000 ORF’s from the *Escherichia coli* genome [5,6]. LASSO probes are designed with two unique annealing arms that are connected through a common nucleotide backbone. In presence of a targeted sequence the arms hybridize to the beginning and the end of the sequence then the sequence is replicated by gap filling and ligation to the ending arm (mentioned herein as ligation arm). The captured DNA sequences can then be reliably cloned in expression systems such as phage, ribosomal display and reporter assays in a massively parallel format.

This report describes the evaluation of key variables involved in the target capture process in library format, specifically: DNA polymerase type, DNA ligase concentration, different LASSO backbone lengths and melting temperature of the arms (**Figure 1**). In summary, target capture process efficiency was improved by increasing the ligase concentration by 10 fold. We tested different DNA polymerases and found that Kapa HiFi polymerase was especially favorable for longer target capture up to 5Kb in size. We also tested the capture efficiency of pools of probes with arms of varying melting temperature in targeting ORFs from *E*.*coli* str. k-12 genome and sequenced the resulting captured libraries. We observed that the pool that had the melting temperature of the extension and ligation arm in the same range of 65-70 C was able to capture homogeneously (MLD of 0.77) 96% of the targeted ORFs. In addition, this pool resulted in 315 fold enrichment of coverage for captured target versus captured non targeted ORF’s. These results begin to codify important standard operating procedures for the critical unit operations involved in library cloning by LASSO probes.

**Figure 1.**
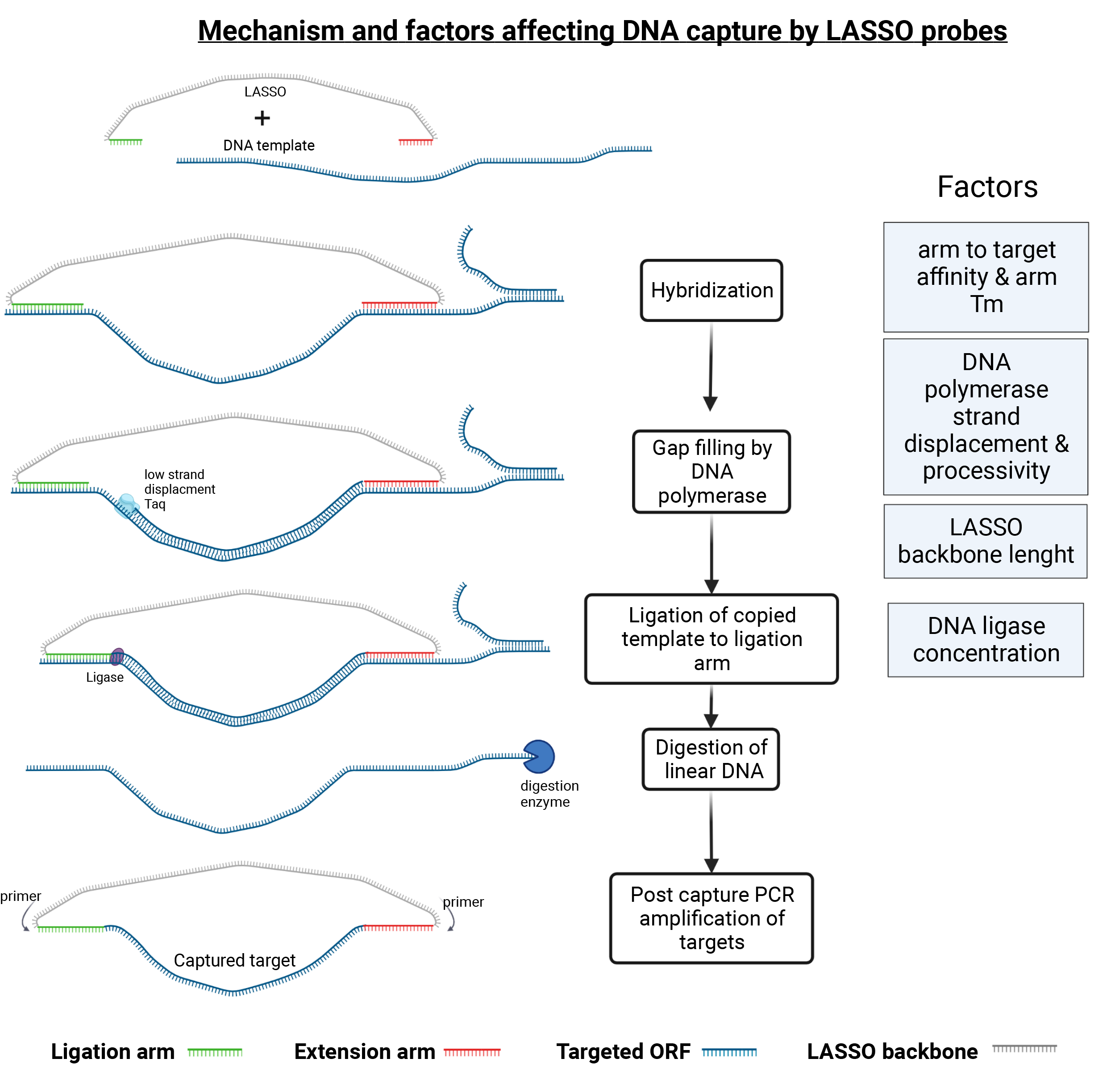

## Materials and methods

### Design of single pre-LASSO probes that target M13mp18 bacteriophage sequences

Pre-LASSO probe pools are short DNA oligo pools (∼160-180 bp) that we first designed in silico and then ordered from Twist Bioscience and used for the assembly of LASSO probes. Pre-LASSO probes have five different regions: primer-annealing site, ligation arm, conserved region, extension arm, primer-annealing site. The ligation and extension arms of the pre-LASSO probes are designed to have the same 5’-3’ orientation of the sequence of the target DNA.

As a positive control, we used the same pre-LASSO probe targeting a 1Kb target capture on the ssDNA of M13mp18 as the one listed by Chkaiban et al. (2021) [7]. It had the Tm of the extension arms ∼ 65°C and the Tm of the ligation arms ∼70°C. Pre-LASSOs targeting a single 3 Kb sequence within the M13mp18 genome were manually designed with Tm of the extension arms ∼ 65°C and 3 different Tm of the ligation arms 65°C, 70°C and 75°C. In addition, pre-LASSO probes targeting single 4 and 5 kb sequences within the single strand M13mp18 DNA were manually designed with Tm of the extension arms ∼ 65°C and the Tm of the ligation arms ∼70°C. The sequences for the above cited pre-LASSO targeting on the M13mp18 genome are listed in **table 1**.

**Table 1.**
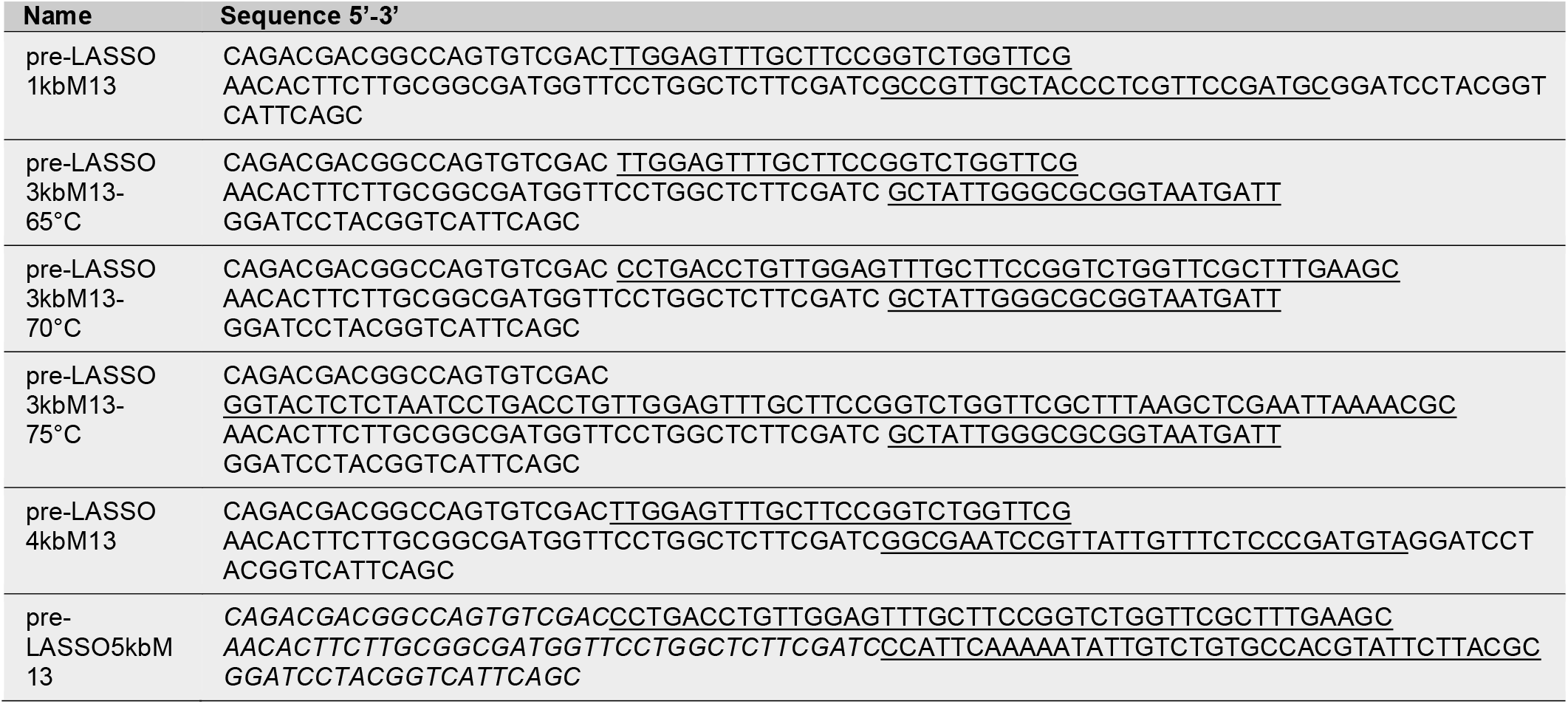
Oligonucleotide list for pre-LASSOs that capture M13mp18 sequences (the ligation and extension arms are underlined)

### Design of different melting arms the pre-LASSO probes pools for an *E. coli* model

We assessed the effect of varying the melting temperature of LASSO probes arms on capture efficiency and specificity by designing probes that targets E.coli ORF’s ranging from 999 bp-2000bp. Specifically, we designed five different pools: a pool that had a 5^o^C lower ligation arm (65-70^o^C) melting temperature with respect to the extension arm (70-75^o^C) (L_Med_E_Hi_); a pool that had a 10^o^C lower ligation arm (60-65^o^C) melting temperature with respect to the extension arm (70-75^o^C) (L_Lo_E_Hi_); a pool that had a 5^o^C lower extension arm (65-70^o^C) melting temperature with respect to the ligation arm (70-75^o^C) (L_Hi_E_Med_); a pool that had a 10^o^C lower extension arm (60-65^o^C) melting temperature with respect to the ligation arm (70-75^o^C) (L_hi_E_Lo_); and a pool that had extension and ligation arm (65-70^o^C) melting temperature in the same range (L_Med_E_Med_). We modified a previously available bio-python algorithm by prolonging the arms until the desired melting temperatures were reached and selected probes that would capture *E*.*coli* ORF targets ranging form 999 bp to 2000 bp [7]. We ran the bio-python algorithm on the *Escherichia coli* str. k-12 substr. mg1655 reference ORFeome found in NCBI (RefSeq: NC_000913.3). The new biopython algorithms as well as the resulting pre-LASSO list of probes can be found in the **supplementary files**.

### Assembly of the LASSO probes

The assembly of the LASSO probes was done using a 350bp backbone according to the protocol described by Chkaiban et al. (2021) for all single LASSOs and LASSO pools [7]. In addition to the assembly with 350bp backbone, to assess the effect of backbone length on capture efficiency we assembled LASSO probes that target 3 Kb sequences in the M13mp18 bacteriophage using a longer 700 bp backbone linker. The 700 bp backbone linker was substituted to the 350bp backbone in the support protocol 1 in the pLASSO plasmid generation listed in Chkaiban et al. (2021) ahead of the LASSO probe assembly protocol [7]. The backbone linker oligonucleotides are listed in **table 2**.

**Table 2.**
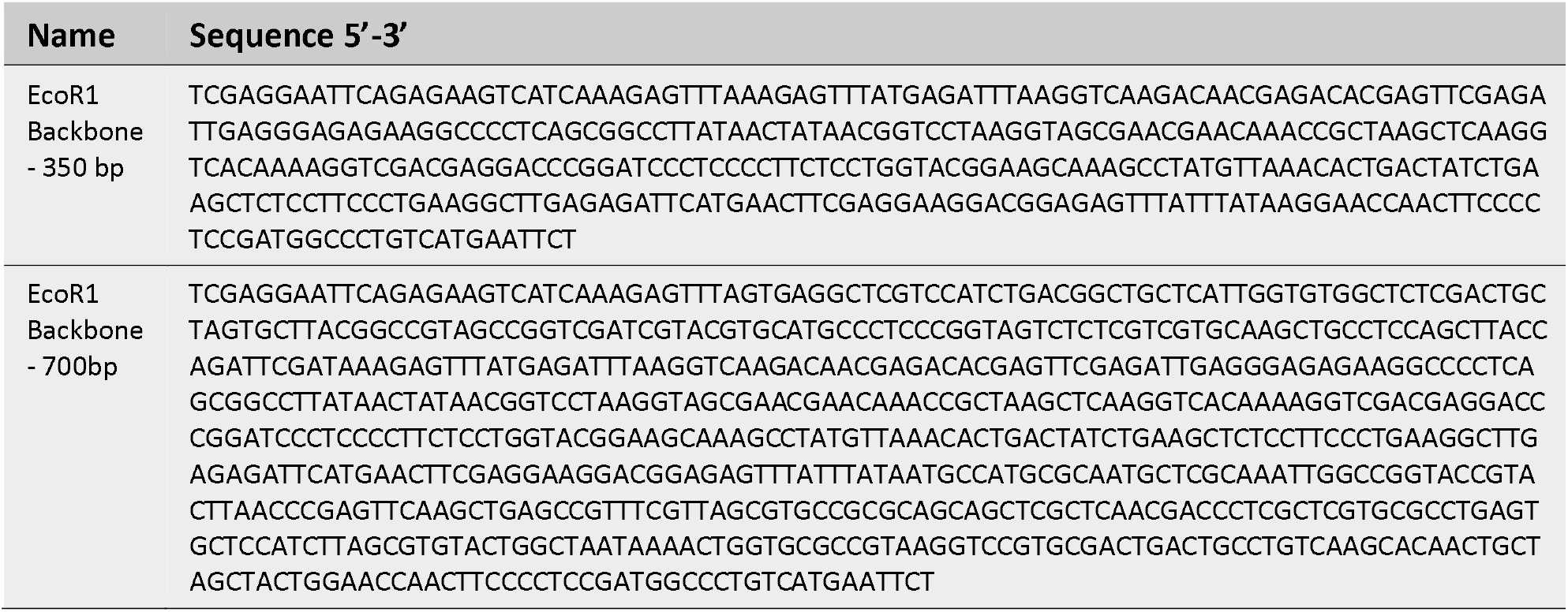
Oligonucleotide list for the backbone linkers

### DNA target capture

In order to optimize capture efficiency, we have tested two different DNA polymerases (Omni Klentaq LA and Kapa HiFi) in the gap filling Mix of the capture step (see **table 3**) with LASSO probes that target 1 Kb and 3 Kb within M13mp18 bacteriophage genome. We also tested 3 ten-fold increases in concentrations of Ampligase DNA Ligase in the gap filling Mix of the capture step with LASSO probes that target 1 Kb on single stranded and double stranded DNA of the M13mp18 bacteriophage (see **table 4**).

**Table 3.** Components of the gap filling Mix with Omni Klentaq LA

| COMPONENT | Amount PER 100 µl<br>Stock |
| --- | --- |
| Omni KlenTaq gap filling mix | 0.5 U ligase |
| PCR grade Water | 75.1 µl |
| 10X Ampligase DNA ligase Buffer | 10 µl |
| 10 mM dNTPs | 0.4 µl |
| Ampligase DNA Ligase (100U/ul) | 0.1 µl |
| Omni Klentaq LA | 0.4 µl |
| NADH | 4 µl |
| Glycerol | 10 µl |

**Table 4.**
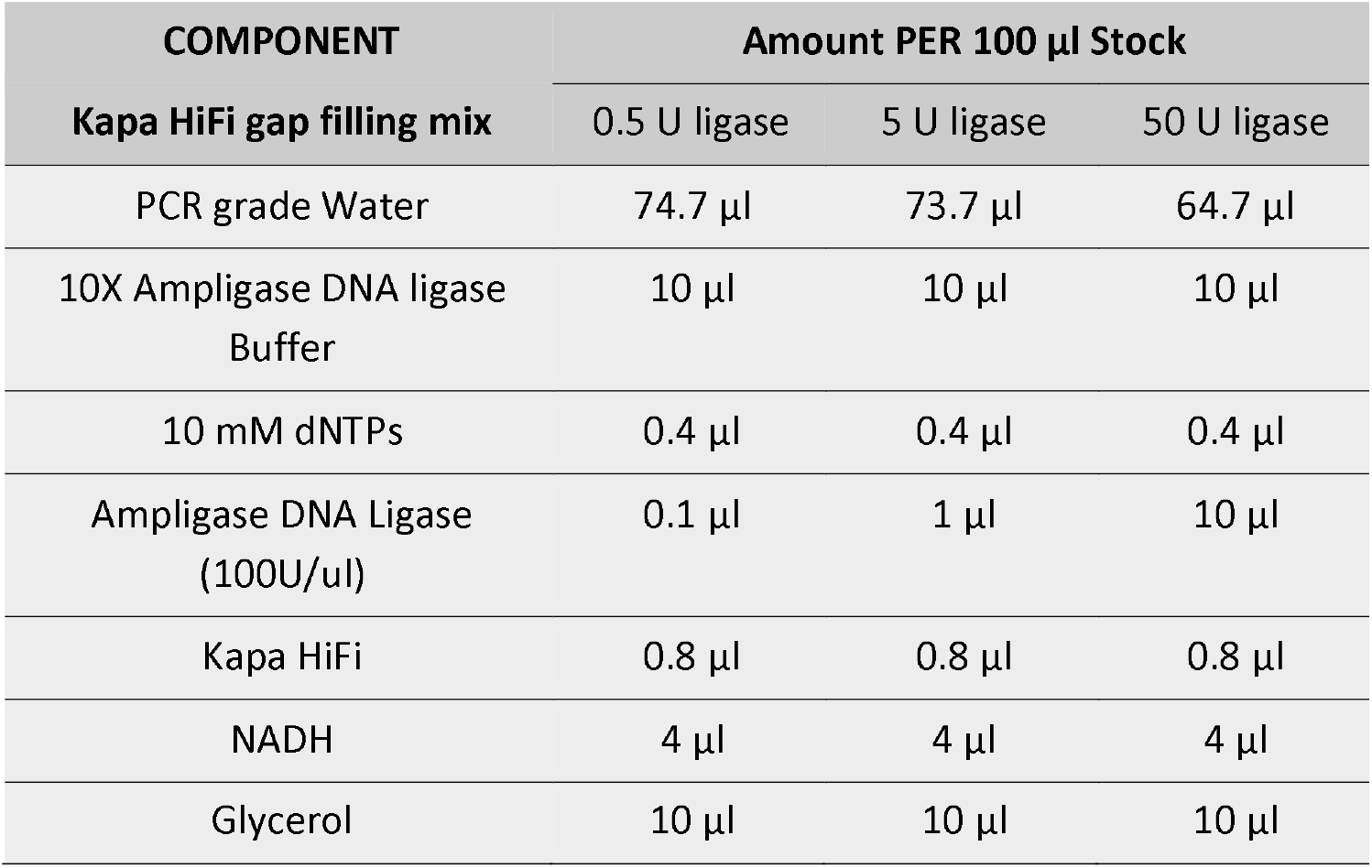
Components of the gap filling Mixes with Kapa HiFi at various concentrations

| COMPONENT | Amount PER 100 µl Stock |  |  |
| --- | --- | --- | --- |
|  | 0.5 U ligase | 5 U ligase | 50 U ligase |
| **Kapa HiFi gap filling mix** |  |  |  |
| PCR grade Water | 74.7 µl | 73.7 µl | 64.7 µl |
| 10X Ampligase DNA ligase Buffer | 10 µl | 10 µl | 10 µl |
| 10 mM dNTPs | 0.4 µl | 0.4 µl | 0.4 µl |
| Ampligase DNA Ligase (100U/ul) | 0.1 µl | 1 µl | 10 µl |
| Kapa HiFi | 0.8 µl | 0.8 µl | 0.8 µl |
| NADH | 4 µl | 4 µl | 4 µl |
| Glycerol | 10 µl | 10 µl | 10 µl |

The Kapa HiFi based gap filling mix and 5 U ligase in the final reaction volume was used for most of the captures, namely: LASSOs targeting 3 Kb sequences within the single strand M13mp18 DNA with 3 different Tm of the ligation arms 65°C, 70°C and 75°C, with 350bp and 700bp backbone linker, LASSOs targeting 4 and 5 Kb sequences within the single strand M13mp18 DNA, and LASSO probes pools that target E. coli DNA and have different melting temperature arms.

The capture was completed with a digestion step after which we performed a post-capture PCR according to the protocol listed in [7]. The primers used in the post capture PCR reaction are listed in **table 5**. We used the total amount of post-capture product measured as fluorescence on gel electrophoresis against known calibration standards as an estimate of the efficiency of the capture reaction.

**Table 5.**
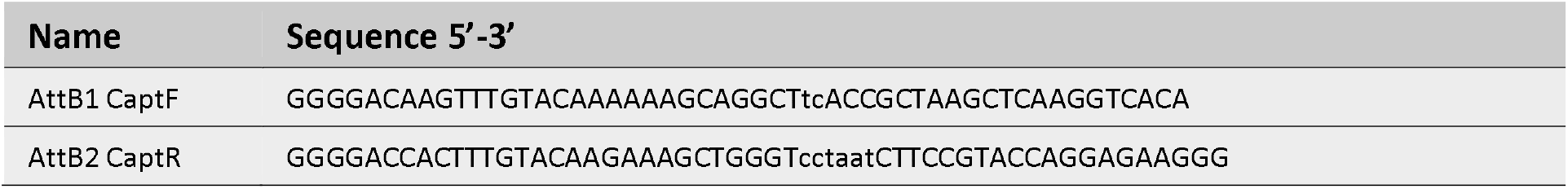
Oligonucleotide list for primers used in the post capture PCR

### Sanger sequencing

We excised the band from the electrophoresis gel showing a 5 Kb captured target band size from the ssDNA of M13mp18 and purified it using Monarch DNA Gel Extraction Kit (#T1020S). We performed a Sanger sequencing on the eluate to confirm the identity of the band.

### DNA preparation and barcoding of pools for Oxford nanopore sequencing

We used the ligation kit SQK-LSK 109 with the PCR barcoding expansion 1–12 EXP-PBC0001 supplied by Oxford Nanopore and followed the respective protocols for DNA sample preparation for sequencing. We primed a R 9.4.1 flow cell with the component supplied in the flow cell priming kit (EXP-FLP002) and loaded 50 fmol after mixing it with loading beads and sequencing buffer supplied with the kits. We ran the sequencing in the MinION Mk1C and set it for real-time data acquisition and basecalling.

### Sequencing data analysis

We aligned the resulting reads founds in fastq files subdivided according to their barcode directly in the MinKNOW app built-in the MinION Mk1C. Each pool was mapped against the ORFeome reference file for *Escherichia coli* str. k-12 substr. mg1655 found in NCBI (RefSeq: NC_000913.3) uploaded locally as a fasta file. The filtering, the statistical analyses and resulting bean plot graph were performed on R software.

### Cloning the captured amplicons pools in the Gateway system

The post capture PCR product pools were bead purified and mixed with the Gateway ‘donor vectors’ (pDONR221) and the BP Clonase enzyme mix (Invitrogen). We purified the BP reaction and used if for electroporation in NEB® 10-beta Electro-competent E. coli (c3020K) to generate cloned libraries. We then extracted the plasmids and digested them with EcoRV restriction enzymes to linearize them and proceeded with end repair and DNA preparation for sequencing with the same ligation and barcoding kit used for the amplicon pools mentioned above (SQK-LSK109 with EXP-PBC001).

## Results

### Effect of DNA polymerase type on capture efficiency

At a single target level, LASSO probes have two arms that bind to flanking DNA regions surrounding an open reading frame (ORF) of interest. This binding event enables the copying of the ORF by a DNA polymerase. Selecting an optimal DNA polymerase that has a good processivity and easily dissociates without displacing the ligation arm after it has completed the replication of the DNA target can greatly improve the efficiency of the capture of a DNA target (see Figure 1). We therefore assessed 2 different DNA polymerases (Omni Klentaq LA and Kapa HiFi) when capturing 1Kb, 3kb and 5kb single targets within ss DNA of the M13mp18 phage genome while all the other components of the gap filling mixes remained the same. The 2 polymerases did not have a significantly different effect on the 1 Kb target capture -estimated in ng of PCR post capture product. While attempting to capture a 3kb target, Kapa HiFi produced a visible capture band most of the time while the Omni Klentaq produced only very weak or no visible bands on the gel as reflected by the large standard deviation for these assays (some gels shown in **supplementary figure 2**). For the 5 Kb target only Kapa Hi Fi was able to generate a visible capture band in the gel (see **supplementary figure 2**). For this higher observed yield in longer target captures we decided to adopt the Kapa HiFi for all the following experiments.

**Figure 2.**
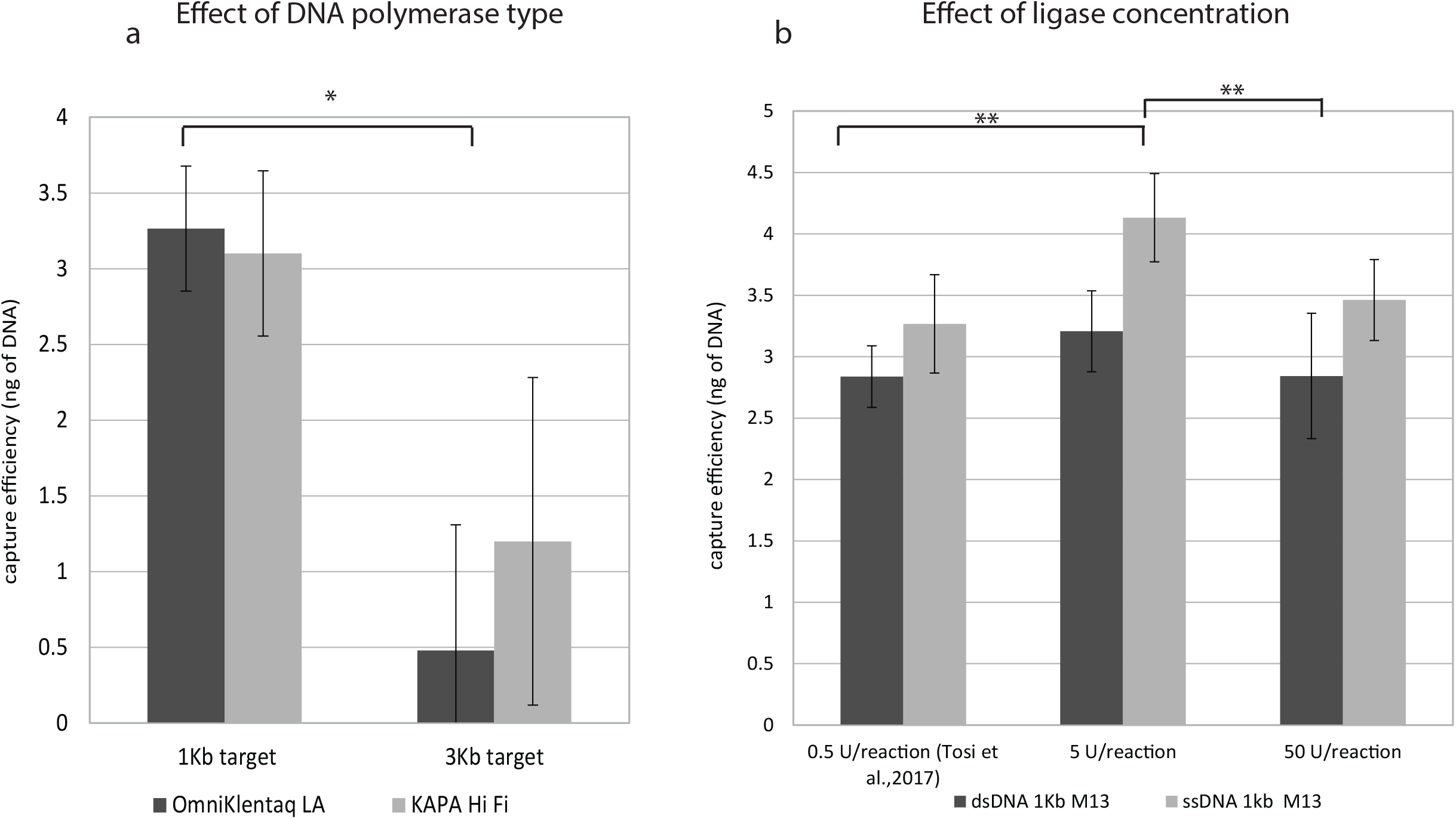

Then, we assessed increasing concentrations of DNA ligase in the gap filling mix (by 10 fold increases) on capture efficiency that we estimate as ng of post capture PCR product measured as fluorescence on gel electrophoresis against known calibration standards. Capture on single strand DNA templates produced higher capture efficiency then when starting with double stranded DNA templates (**Figure 2b**). In addition we noted that among the 3 conditions tested, the 5 unit of DNA ligase/in the 20 µl reaction volume was the optimal concentration in terms of capture efficiency (**Figure 2.b**).

### Effect of different LASSO backbone lengths and melting temperature ligation arms on capture efficiency of single targets

LASSO probes represent an improvement over probes that have shorter backbone (∼100nt) such as molecular inversion probes because the longer backbone that links the arms in the LASSO probes allow for the capture of longer DNA targets [8]. Here, we further investigate the effect of extending the length of the backbone on capture efficiency. In addition, LASSOs arms have so far been designed with the ligation arm having a melting temperature 5°C higher than the extension arm (that is ∼65°C) because it was hypothesized to increase the strength of the hybridization of the ligation arm to DNA target. We have therefore tested the effect of backbone length in combination with various melting temperature of the ligation arm on capture efficiency. We assembled 6 LASSO probes having 3 progressively higher Tm ligation arms for 2 different length backbones (350 and 700 bp). The 6 LASSO were designed to target the same 3Kb region on ssDNA of M13mp18 phage (see **supplementary figure 1**). We observed that LASSOs with the shorter 350 bp backbone performed better than with longer backbone 700bp, which was especially evident for 1 Kb targets (**Figure 3 a** and **b**). Hence, we adopted the 350 bp backbone for LASSO probes assembly as best practice for subsequent experiments. **Figures 3 a** and **b** also show the effect of melting temperature of the ligation arm on capture efficiency. The highest capture efficiency was obtained when using a ligation arm of 70°C melting temperature.

**Figure 3.**
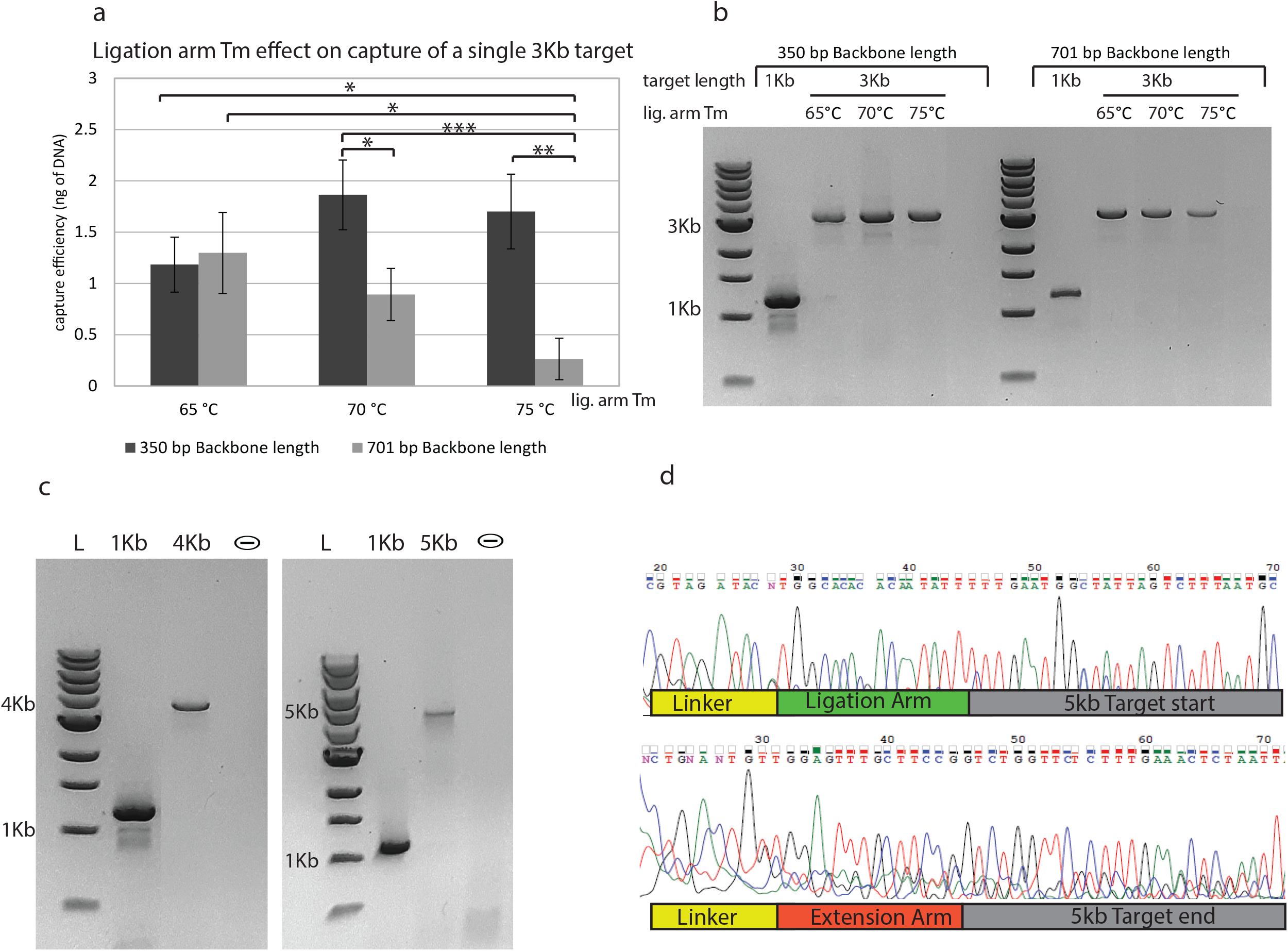

### 4 and 5 Kb long single targets capture

The presence of genetic elements that encompass several Kb in size creates the need for technologies that can capture pools of full-length DNA targets. In order to test the capability of the LASSO technology in capturing long DNA targets, we designed pre-LASSO probes that target single 4 and 5 Kb sequences on M13mp18 genomic ssDNA with Tm of the extension arms ∼ 65°C and the Tm of the ligation arms ∼70°C. When running the post capture PCR product on an electrophoresis gel we were able to detect bands at around 4kb and 5kb that indicate successful capture of our targeted sequences (**Figure 3.c**). Furthermore, we were able to corroborate the identity of the 5kb band by Sanger sequencing it after excising and purifying it from the gel. The 2 chromatograms, obtained by sequencing with forward and reverse post capture PCR primers, showed at the ends the presence of a sequence that mapped with ligation (in green) and extension (in red) arms as per the design of the probe (**Figure 3.d**) followed by the rest of the targeted sequence indicating that the target was captured in its full length.

### Testing capture efficiency of LASSO pools with different melting temperatures arms

In an effort to establish more accurate and improved parameters for the design of pre-LASSOs we tested LASSO pools of varying melting temperature (Tm) of ligation and extension arms when capturing targets within the *Escherichia coli* ORFeome ranging from 999 bp-2000bp (**Figure 4 .a**). **Figure 4. c** shows the distribution of the potential targets of the designed LASSOs by length into bins of 50 bp incrementally. Most of the LASSOs target sequences are in the range of 1000 to 1400 bp in size. The algorithm produced 128 to 807 LASSOs out of the 4140 ORF per pool. Running the post capture product on a gel of the various captured pools showed a smear for each pool in the expected size range (**Figure 4. b**). The smear were more pronounced in the range of 1000 to 1400 bp which is in accordance to the size distribution initially produced by the algorithm.

**Figure 4.**
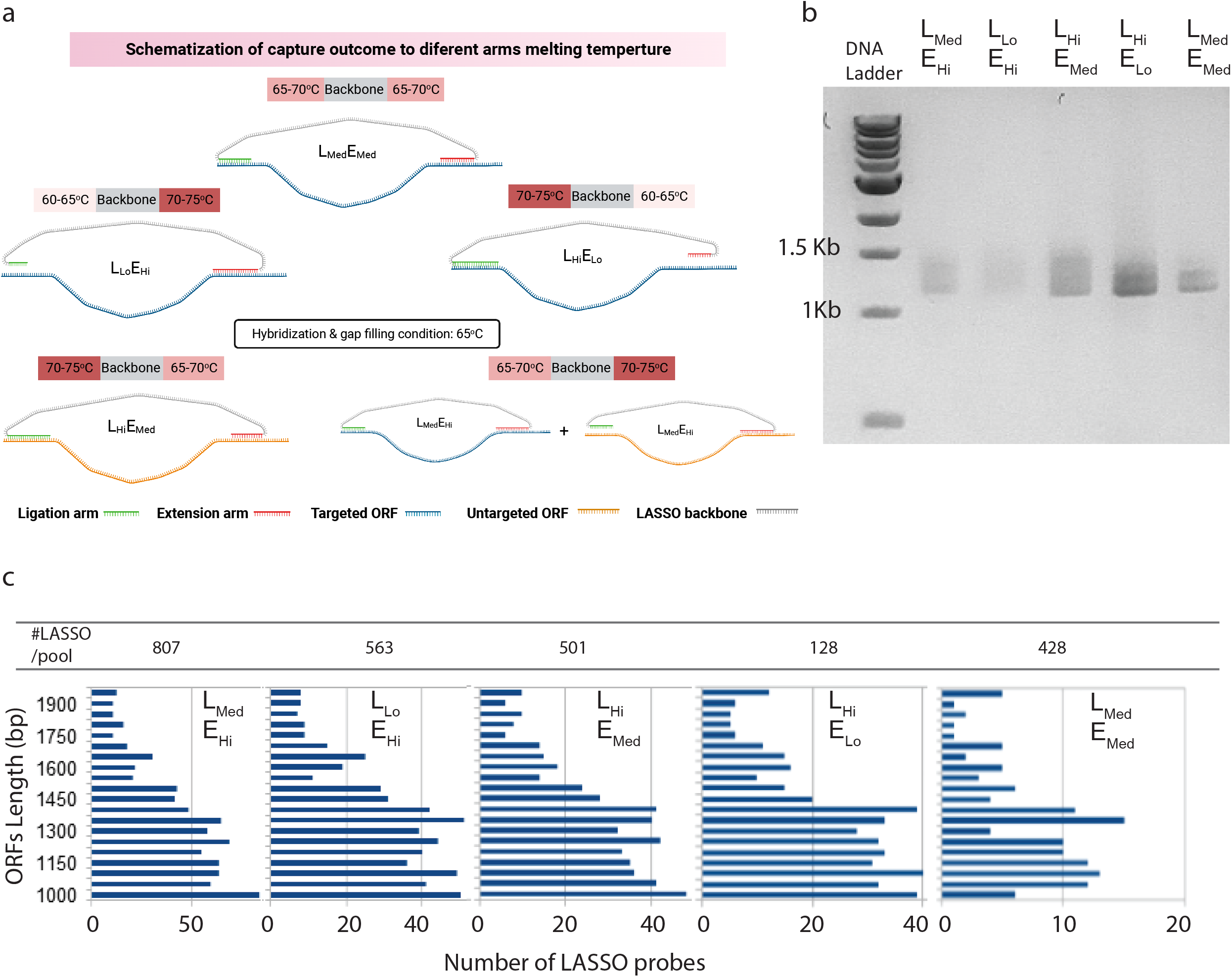

The post capture amplicons were obtained by amplification by using attB tailed primers thus sequenced with MinION Mk1C. Using R software we calculated the depth of coverage for each target and plotted it for both the pools of captured amplicons (**Figure 5.a**) and the pools of the amplicons transformed into pDNOR 221 plasmids (**Figure 5.b**). With respect to the pools of amplicon targets, the highest coverage on average was obtained for the targeted ORFs captured with the L_Hi_E_Med_ pool that had the melting temperature of the extension arm in the range of 65-70°C and ligation arm in 70-75°C, whereas the most homogeneous distribution was observed in the L_Med_E_Med_ because it yielded the lowest mean log deviation (MLD) of 0.77 in comparison to 2.90, 3.73, 2.06, 3.24 of the L_Med_E_Hi_, L_Lo_E_Hi_, L_Hi_E_Med_, L_Hi_E_Lo_ pools respectively. We used the mean log deviation (MLD) as an indicator dis-proportionality in the coverage of targeted of ORF. When we filtered and computed the coverage of non-targeted ORFs for each pool, we observed that the median coverage was between 0.91 for L_Med_E_Med_ and 63.99, for L_Hi_E_Med_ (**Figure 5.a**). This shows the low specificity for probes (L_Hi_E_Med_ pool) having melting arm temperature of ligation ∼5°C higher than the extension (65-70°C) while the highest specificity was for the pool L_Med_E_Med_ that had extension and ligation arm melting temperature in the same range (65-70°C), recorded as lowest coverage for untargeted sequences. In addition, we observed that at a cutoff of three times the median non-target coverage, 96% of the targeted ORFs were successfully captured by L_Med_E_Med_ indicating the higher capture efficiency for that pool as shown in **Figure 5. c**. In addition, targets were 57, 0.9, 7, 4 and 315 fold enriched with respect to non-targeted ORF’s for each of L_Med_E_Hi_, L_Lo_E_Hi_, L_Hi_E_Med_, L_Hi_E_Lo_ and L_Med_E_Med_ pools respectively.

**Figure 5.**
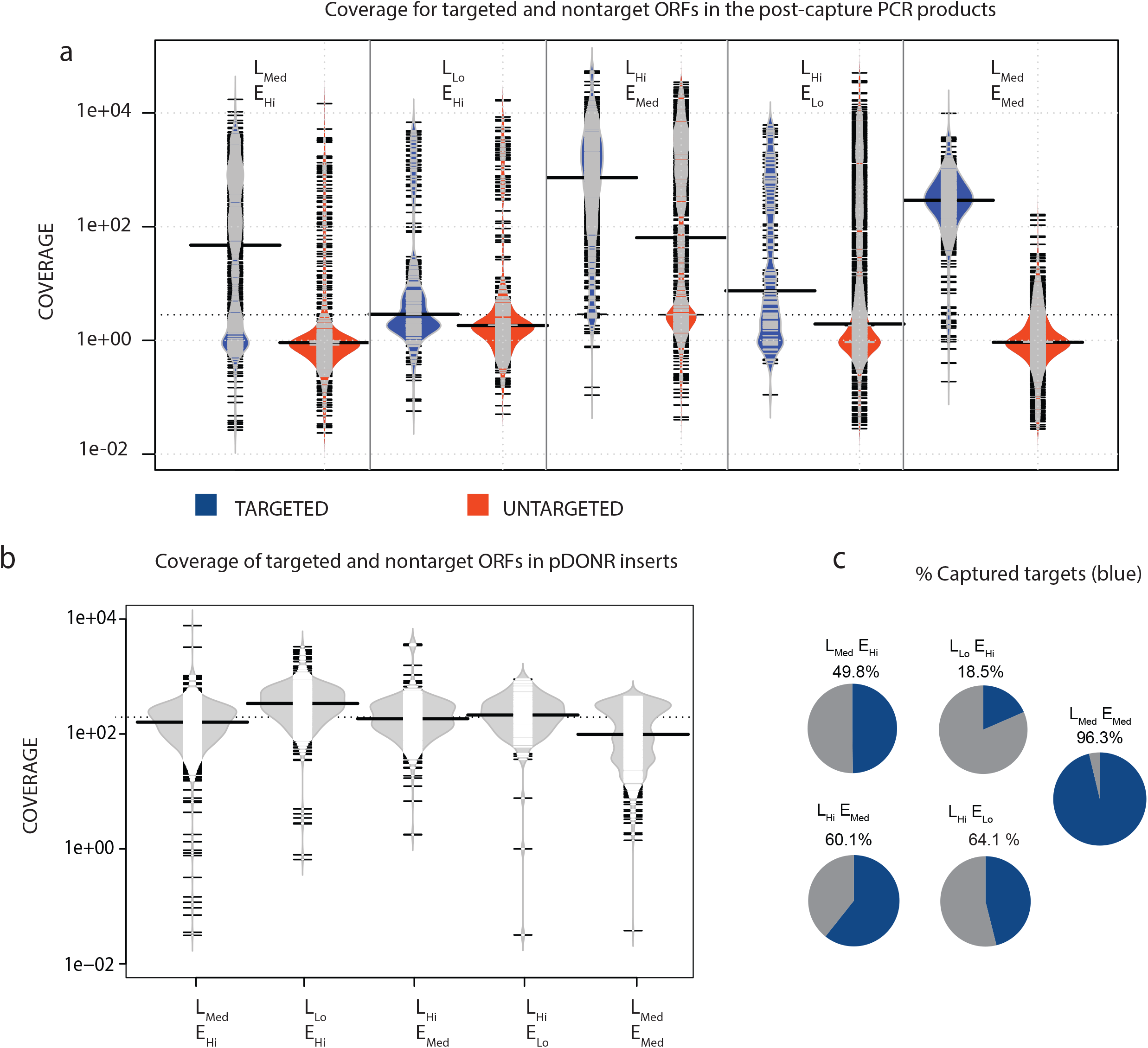

In order to test the applicability of the captured libraries for cloning purposes we tested the possibility of transferring our captured pools into a widely used cloning system, the pDONR221 from the Gateway technology. The amplicons were cloned into pDONR221 via BP recombination reaction and used for transforming electrocompetent *Escherichia coli*. The antibiotic resistant colonies were selected from agar plates and the extracted plasmids were sequenced with MinION Mk1C. We have observed that the coverage of the cloned libraries did not represent the frequency distribution of their respective un-cloned counterpart and the coverage of targeted and cloned ORFs displayed narrower distribution (**Figure 5. b**) with similar median coverage, respectively 161, 339, 184, 214 and 99 for each of pDONR221 cloned L_Med_E_Hi_, L_Lo_E_Hi_, L_Hi_E_Med_, L_Hi_E_Lo_ and L_Med_E_Med_ pools.

## Discussion

LASSO probes are in evaluation as a tool to unravel DNA libraries through cloning and functional expression analysis. This report characterized the impact of process controls for the gap filling reaction components together and design of the probe arms and verified the impact on capture efficiency. Tosi and colleagues (2022) recently have re-engineered the LASSO assembly methodology that was previously fusion PCR-based into a Cre-recombinase mediated assembly technique [6]. This resulted in better quality LASSO probes which had an impact on the quality of a captured pool. In fact, the sequencing of the captured targeted ORF’s showed an improvement in enrichment of 8 times higher for the CRE-recombinase mediated assembled LASSOs with respect to the fusion assembled LASSOs. CRE-mediated assembly was thus used in this study to target and clone thousands of ORF’s using LASSOs.

LASSO’s DNA target capture mechanisms are similar to that of inversion probes (MIPs) in that they both capture by gap filling and ligation. MIPs are limited to a maximum ∼150 bp target length because of the shortness of the backbone that does not allow for the capture of long targets [8, 9]. During the gap filling step as the target sequence is replicating it becomes double stranded because of the repulsion of ionized phosphodiester bonds on the opposite strands the dsDNA target stiffens (persistence length 50nm (or 150 bp) in 0.1 M NaCl aqueous solution) [11]. In a previous attempt, longer capture of 500∼600bp targets were achieved by extending the linker of capturing probes, however the method relied on individual PCR to extend the length of the initial probe which is not easily scalable [11, 12]. The multiplexed assembly combined with the further increase in probe length (∼450 nt) is the major novelty of the LASSO probes that allows for longer captures, up to several Kb. In this study, in an effort to explore the effect of longer backbones (previously backbones length∼ 350bp) we have attempted to capture with LASSO having 700 bp long backbone. Our results show that exceeding a certain limit in the backbone linker was counter-productive in terms of capturing efficiency. This is in accordance to previous simulation studies performed on LASSO that showed that increasing backbone linker over a certain length could enhance unzippping forces within dsDNA [14,15].

MIPs capturing ability was improved over time by: increasing the amount of time allowed for hybridization reactions, increasing the amounts of reactants required to ensure adequate generation of circles, and careful adjusting dNTP concentrations to minimize the strand displacement activity of the polymerase [16,17]. A critical reagent in the capture process is the DNA polymerase that extends the 3’ end starting from the extension arm thus copying the target sequence till the ligation arm. Upon arrival at the ligation arm end, the DNA polymerase has to dissociate to allow for the DNA ligase to ligate the 5’ end of the ligation arm with the newly synthesized DNA. For this reason, it is important to select a DNA polymerase that has a low strand displacement ability so it can dissociate when it reaches the ligation arm. The stoffel fragment of the AmpliTaq DNA polymerase (Applied Biosystems) is commonly used for gap filling when performing multiplex target capture (size ∼40 nt) with padlock probes and MIP for its lack of 5’→3’ exonuclease activity and its low strand displacement activity [18,19]. The stoffel fragment was found to be better than Phusion High-Fidelity DNA Polymerase (Thermo Fisher Scientific) in DNA gap filling [17]. When capturing ORF targets (size 400 up to 4Kb) with LASSOs researchers have used Omni Klentaq LA (DNA polymerase technologies) because of its low strand displacement activity [5,6,20,21]. A number of authors have reported good processivity and low introduction of bias by Kapa HiFi DNA polymerase [22–24]. We tested this enzyme in the gap filling mix and found that it over-performed Omni Klentaq LA for longer target captures, which may be due to similar if not weaker strand displacement ability of Kapa HiFi with respect to Omni Klentaq LA.

We observed that increasing the concentration of DNA ligase by 10 folds resulted in an improved target capture efficiency (from 0.5 to 5 U in 20ul of the reaction volume). This is an agreement with a previous study where the authors have observed an improved molecular inversion probes (MIPs) capture efficiency for a human exon library when increasing ligase concentrations (0.25 U → 5 U in a ∼20 µL reaction volume) [25]. This suggests that this higher concentration of ligase in combination with that of DNA polymerase (0.16 U/µl.) maximizes the number of the successfully closed circles. Using this new gap filling mix composition, we were able to capture a longer target (5Kb) with respect to previously reported ∼4 Kb target capture [5,6].

The main challenge of the LASSO capture is to design a pool of probes that can capture their targets with similar efficiencies so that in the final captured library all the targets are represented with the similar frequency. One of the main variables when designing the probes is the Tm of the arms. To minimizes a possible polymerase displacement of the hybridized arm to the template some authors proposed to design probes with 5°C higher meting temperature in the ligation arms with respect to the extension arm, suggesting that it would stabilizing the ligation arm at the 5’ ligation site [18,26]. This design was also previously adopted for both MIPs and LASSO probes [5,19]. We have designed pools of LASSOs that have varying Tm arms and tested their efficiency in capturing a multiplex long *Escherichia coli* ORF ranging from 1kb to 2kb. We observed that the subpool with ligation arm 5 C higher than that of the extension arm 65-70°C named L_Hi_E_Med_ yielded the highest median coverage for the captured targets. However, it resulted in a non-homogeneous capture with many over and under-represented ORFs and a relatively high nonspecific capture. The best capture achieved in this work following optimization in DNA ligase concentration, DNA polymerase type, and Tm of LASSO arms scored a 315-fold enrichment of targeted versus non-targeted genomic regions while previous LASSO pool capture studies have reached 60-fold enrichment which represents a 5-fold improvement [5,6]. In addition, our data showed that the best capture uniformity, in terms of probes representation, highest target enrichment and specificity and almost complete capture of all targets was obtained with probes designed with equal melting arm temperature in the 65-70°C range. With the hybridization and gap filling conditions set to occur at 65°C, a temperature optimal for proper functioning of DNA ligase and polymerase, we hypothesize that a ligation arm that melt at higher temperature, as it is for the case of L_Hi_E_Med_ pool, may lead to unspecific annealing of the ligation arm to unwanted targets, which penalizes the captures in specificity and uniformity (**Figure 4 a**).

## FUTURE PERSPECTIVE

High-throughput phenotyping technologies require large DNA libraries to represent functional genetic elements that can be several Kb in size. Despite the improvement in silicon-based DNA synthesis, the DNA maximum length for large pools of DNA sequences is still limited to 300 bp to date. Therefore, there is an unmet need for the production of large DNA libraries spanning the length of full genes or long DNA regions at low cost.

This work has focused on optimizing LASSO capture by testing different types of DNA polymerases and concentrations of the DNA ligase in the gap filling mix and by tuning backbone length and melting temperature of the arms in both single target and multiplex target capture experiments. The amendments resulted in longer (up to 5Kb), more specific and homogeneous captures. This improvement will be essential for the applicability of LASSO for the construction of full length ORF or DNA regions for downstream high throughput functional studies. We also envision that LASSO target capture can effectively supplement third generation sequencers by honing on Kb size DNA regions of interest.

## SUMMARY POINTS

- LASSO are Long-Adapter Single-Strand Oligonucleotide that can massively clone long DNA regions of interest in parallel.
- LASSO DNA captures do not benefit from increasing the length of the backbone linking the arms over 350bp
- We improved the single target capture process efficiency by increasing the ligase concentration by 10 fold.
- Kapa HiFi polymerase was especially favorable for longer target captures.
- LASSO can capture targets 5Kb in size.
- L_Med_E_Med_ LASSO pool that has a melting temperature of the extension and ligation arm of LASSO in the same range of 65-70°C was able to capture homogeneously (MLD of 0.77) 96 % of the targeted ORFs.
- L_Med_E_Med_ pool resulted in 315 fold enrichment of coverage for captured target versus captured non targeted ORF’s.

## Abbreviations

Tm: melting temperature
MLD: mean log deviation
LASSO: Long-Adapter Single-Strand Oligonucleotide

## SUPPLEMENTARY FILES

Biopython algorithms, pre-LASSO list of probes, supplementary figures

