## Supplementary figures and images for "Capture Efficiency Of Long-Adapter Single-Strand Oligonucleotide Probe Libraries"

### supplementary figure 1

#### LASSOs targeting a single 3Kb target on M13 mp18 genome

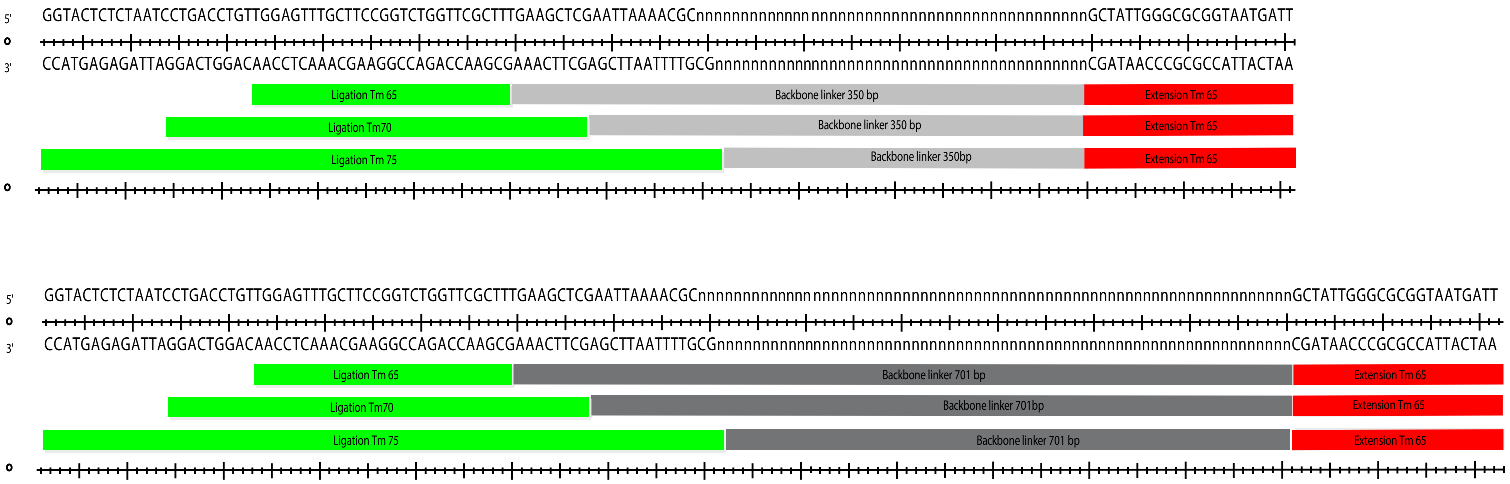

### supplementary figure 2

#### Long single target capture essay: Omni Klentaq vs Kapa Hi Fi

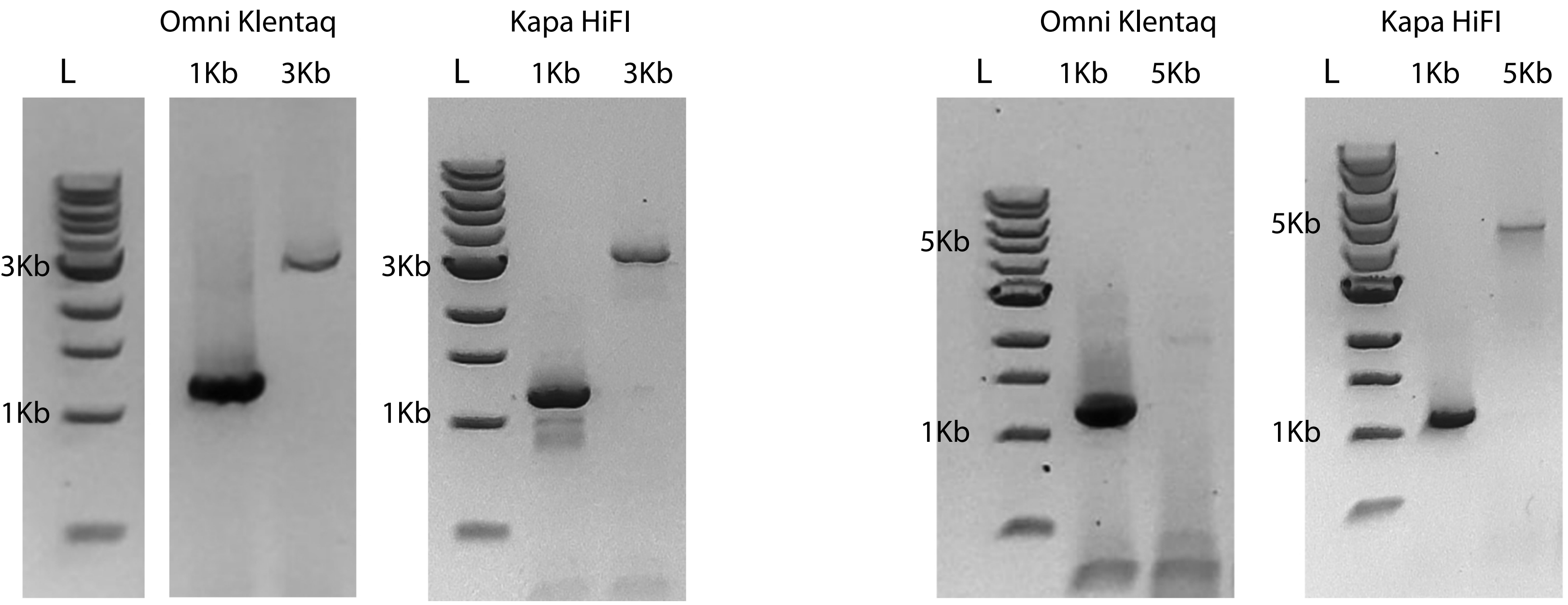
