## supplementary files for "Capture Efficiency Of Long-Adapter Single-Strand Oligonucleotide Probe Libraries"

LMed EMed biopython script

-----------------------------------

from Bio import SeqIO

import random

from Bio.SeqUtils.MeltingTemp import Tm_staluc

r = {}

for i,seq_record in enumerate(SeqIO.parse("k12/k12.ffn","fasta")):

print i

s = str(seq_record.seq)

ls = len(s)

if ls < 200:

continue

r[seq_record.id] = []

r[seq_record.id].append(s)

r[seq_record.id].append(seq_record.description)

r[seq_record.id].append(str(ls))

failed = False

v = 32

while 1:

if v >= len(s) - 1:

failed = True

break

while s[v] not in ['C','G']:

if v >= len(s) - 1:

failed = True

break

v += 1

if failed:

break

start = s[:v]

stop = s[-(100-len(start)+3):-3]

tmstart = Tm_staluc(start)

tmstop = Tm_staluc(stop)

if len(start) > 60:

failed = True

break

if tmstart < 65.0:

v += 1

continue

if tmstop < 65.0:

print "tmstop < 60.0"

failed = True

break

if tmstop > 70.0:

v += 1

continue

if tmstart > 70.0:

print "tmstart > 65.0"

failed = True

break

if tmstop < tmstart:

v += 1

continue

break

if failed:

r[seq_record.id].append('FAILED')

continue

r[seq_record.id].append(start)

r[seq_record.id].append(str(len(start)))

r[seq_record.id].append(stop)

r[seq_record.id].append(str(len(stop)))

r[seq_record.id].append(str(len(start)+len(stop)))

r[seq_record.id].append(str(Tm_staluc(start)))

r[seq_record.id].append(str(Tm_staluc(stop)))

r[seq_record.id].append('cAgACGACGGCCAGTgtcgac'+start)

r[seq_record.id].append('cAgACGACGGCCAGTgtcgac'+start+'AACACTTCTTGCGGCGATGGTTCCTGGCTCTTCGATC')

r[seq_record.id].append('cAgACGACGGCCAGTgtcgac'+start+'AACACTTCTTGCGGCGATGGTTCCTGGCTCTTCGATC'+stop)

r[seq_record.id].append('cAgACGACGGCCAGTgtcgac'+start+'AACACTTCTTGCGGCGATGGTTCCTGGCTCTTCGATC'+stop+'GGATCCTACggtcATtCAGC')

toadd = 180 - len('cAgACGACGGCCAGTgtcgac'+start+'AACACTTCTTGCGGCGATGGTTCCTGGCTCTTCGATC'+stop+'GGATCCTACggtcATtCAGC')

sta = "TGA"

c = {0:'A', 1:'T', 2:'C', 3:'G'}

while "TGA" in sta or "TAA" in sta or "TAG" in sta:

sta = ""

for l in range(toadd):

sta += c[random.randint(0,3)]

r[seq_record.id].append('cAgACGACGGCCAGTgtcgac'+start+'AACACTTCTTGCGGCGATGGTTCCTGGCTCTTCGATC'+stop+sta+'GGATCCTACggtcATtCAGC')

with open("out.ligation~exten.txt","w") as out:

for v in r.values():

out.write( "\t".join(v) +"\n")

-----------------------------------------

L Lo E Med biopython script

----------------------------------------

from Bio import SeqIO

import random

from Bio.SeqUtils.MeltingTemp import Tm_staluc

r = {}

for i,seq_record in enumerate(SeqIO.parse("k12/k12.ffn","fasta")):

print i

s = str(seq_record.seq)

ls = len(s)

if ls < 200:

continue

r[seq_record.id] = []

r[seq_record.id].append(s)

r[seq_record.id].append(seq_record.description)

r[seq_record.id].append(str(ls))

failed = False

v = 32

while 1:

if v >= len(s) - 1:

failed = True

break

while s[v] not in ['C','G']:

if v >= len(s) - 1:

failed = True

break

v += 1

if failed:

break

start = s[:v]

stop = s[-(100-len(start)+3):-3]

tmstart = Tm_staluc(start)

tmstop = Tm_staluc(stop)

if len(start) > 60:

failed = True

break

if tmstart < 60.0:

v += 1

continue

if tmstop < 70.0:

print "tmstop < 60.0"

failed = True

break

if tmstop > 75.0:

v += 1

continue

if tmstart > 65.0:

print "tmstart > 65.0"

failed = True

break

if tmstop < tmstart:

v += 1

continue

break

if failed:

r[seq_record.id].append('FAILED')

continue

r[seq_record.id].append(start)

r[seq_record.id].append(str(len(start)))

r[seq_record.id].append(stop)

r[seq_record.id].append(str(len(stop)))

r[seq_record.id].append(str(len(start)+len(stop)))

r[seq_record.id].append(str(Tm_staluc(start)))

r[seq_record.id].append(str(Tm_staluc(stop)))

r[seq_record.id].append('cAgACGACGGCCAGTgtcgac'+start)

r[seq_record.id].append('cAgACGACGGCCAGTgtcgac'+start+'AACACTTCTTGCGGCGATGGTTCCTGGCTCTTCGATC')

r[seq_record.id].append('cAgACGACGGCCAGTgtcgac'+start+'AACACTTCTTGCGGCGATGGTTCCTGGCTCTTCGATC'+stop)

r[seq_record.id].append('cAgACGACGGCCAGTgtcgac'+start+'AACACTTCTTGCGGCGATGGTTCCTGGCTCTTCGATC'+stop+'GGATCCTACggtcATtCAGC')

toadd = 180 - len('cAgACGACGGCCAGTgtcgac'+start+'AACACTTCTTGCGGCGATGGTTCCTGGCTCTTCGATC'+stop+'GGATCCTACggtcATtCAGC')

sta = "TGA"

c = {0:'A', 1:'T', 2:'C', 3:'G'}

while "TGA" in sta or "TAA" in sta or "TAG" in sta:

sta = ""

for l in range(toadd):

sta += c[random.randint(0,3)]

r[seq_record.id].append('cAgACGACGGCCAGTgtcgac'+start+'AACACTTCTTGCGGCGATGGTTCCTGGCTCTTCGATC'+stop+sta+'GGATCCTACggtcATtCAGC')

with open("out.ligation.low.txt","w") as out:

for v in r.values():

out.write( "\t".join(v) +"\n")

--------------------------------

L Hi E Lo bioppython script

--------------------------------

from Bio import SeqIO

import random

from Bio.SeqUtils.MeltingTemp import Tm_staluc

r = {}

for i,seq_record in enumerate(SeqIO.parse("k12/k12.ffn","fasta")):

print i

s = str(seq_record.seq)

ls = len(s)

if ls < 200:

continue

r[seq_record.id] = []

r[seq_record.id].append(s)

r[seq_record.id].append(seq_record.description)

r[seq_record.id].append(str(ls))

failed = False

v = 32

while 1:

if v >= len(s) - 1:

failed = True

break

while s[v] not in ['C','G']:

if v >= len(s) - 1:

failed = True

break

v += 1

if failed:

break

start = s[:v]

stop = s[-(100-len(start)+3):-3]

tmstart = Tm_staluc(start)

tmstop = Tm_staluc(stop)

if len(start) > 60:

failed = True

break

if tmstart < 70.0:

v += 1

continue

if tmstop < 60.0:

print "tmstop < 60.0"

failed = True

break

if tmstop > 65.0:

v += 1

continue

if tmstart > 75.0:

print "tmstart > 75.0"

failed = True

break

if tmstop > tmstart:

v += 1

continue

break

if failed:

r[seq_record.id].append('FAILED')

continue

r[seq_record.id].append(start)

r[seq_record.id].append(str(len(start)))

r[seq_record.id].append(stop)

r[seq_record.id].append(str(len(stop)))

r[seq_record.id].append(str(len(start)+len(stop)))

r[seq_record.id].append(str(Tm_staluc(start)))

r[seq_record.id].append(str(Tm_staluc(stop)))

r[seq_record.id].append('cAgACGACGGCCAGTgtcgac'+start)

r[seq_record.id].append('cAgACGACGGCCAGTgtcgac'+start+'AACACTTCTTGCGGCGATGGTTCCTGGCTCTTCGATC')

r[seq_record.id].append('cAgACGACGGCCAGTgtcgac'+start+'AACACTTCTTGCGGCGATGGTTCCTGGCTCTTCGATC'+stop)

r[seq_record.id].append('cAgACGACGGCCAGTgtcgac'+start+'AACACTTCTTGCGGCGATGGTTCCTGGCTCTTCGATC'+stop+'GGATCCTACggtcATtCAGC')

toadd = 180 - len('cAgACGACGGCCAGTgtcgac'+start+'AACACTTCTTGCGGCGATGGTTCCTGGCTCTTCGATC'+stop+'GGATCCTACggtcATtCAGC')

sta = "TGA"

c = {0:'A', 1:'T', 2:'C', 3:'G'}

while "TGA" in sta or "TAA" in sta or "TAG" in sta:

sta = ""

for l in range(toadd):

sta += c[random.randint(0,3)]

r[seq_record.id].append('cAgACGACGGCCAGTgtcgac'+start+'AACACTTCTTGCGGCGATGGTTCCTGGCTCTTCGATC'+stop+sta+'GGATCCTACggtcATtCAGC')

with open("out.ligation.hi.txt","w") as out:

for v in r.values():

out.write( "\t".join(v) +"\n")

--------------------------------

L Med E Lo bioppython script

-------------------------------

from Bio import SeqIO

import random

from Bio.SeqUtils.MeltingTemp import Tm_staluc

r = {}

for i,seq_record in enumerate(SeqIO.parse("k12/k12.ffn","fasta")):

print i

s = str(seq_record.seq)

ls = len(s)

if ls < 200:

continue

r[seq_record.id] = []

r[seq_record.id].append(s)

r[seq_record.id].append(seq_record.description)

r[seq_record.id].append(str(ls))

failed = False

v = 32

while 1:

if v >= len(s) - 1:

failed = True

break

while s[v] not in ['C','G']:

if v >= len(s) - 1:

failed = True

break

v += 1

if failed:

break

start = s[:v]

stop = s[-(100-len(start)+3):-3]

tmstart = Tm_staluc(start)

tmstop = Tm_staluc(stop)

if len(start) > 60:

failed = True

break

if tmstart < 70.0:

v += 1

continue

if tmstop < 65.0:

print "tmstop < 60.0"

failed = True

break

if tmstop > 70.0:

v += 1

continue

if tmstart > 75.0:

print "tmstart > 75.0"

failed = True

break

if tmstop > tmstart:

v += 1

continue

break

if failed:

r[seq_record.id].append('FAILED')

continue

r[seq_record.id].append(start)

r[seq_record.id].append(str(len(start)))

r[seq_record.id].append(stop)

r[seq_record.id].append(str(len(stop)))

r[seq_record.id].append(str(len(start)+len(stop)))

r[seq_record.id].append(str(Tm_staluc(start)))

r[seq_record.id].append(str(Tm_staluc(stop)))

r[seq_record.id].append('cAgACGACGGCCAGTgtcgac'+start)

r[seq_record.id].append('cAgACGACGGCCAGTgtcgac'+start+'AACACTTCTTGCGGCGATGGTTCCTGGCTCTTCGATC')

r[seq_record.id].append('cAgACGACGGCCAGTgtcgac'+start+'AACACTTCTTGCGGCGATGGTTCCTGGCTCTTCGATC'+stop)

r[seq_record.id].append('cAgACGACGGCCAGTgtcgac'+start+'AACACTTCTTGCGGCGATGGTTCCTGGCTCTTCGATC'+stop+'GGATCCTACggtcATtCAGC')

toadd = 180 - len('cAgACGACGGCCAGTgtcgac'+start+'AACACTTCTTGCGGCGATGGTTCCTGGCTCTTCGATC'+stop+'GGATCCTACggtcATtCAGC')

sta = "TGA"

c = {0:'A', 1:'T', 2:'C', 3:'G'}

while "TGA" in sta or "TAA" in sta or "TAG" in sta:

sta = ""

for l in range(toadd):

sta += c[random.randint(0,3)]

r[seq_record.id].append('cAgACGACGGCCAGTgtcgac'+start+'AACACTTCTTGCGGCGATGGTTCCTGGCTCTTCGATC'+stop+sta+'GGATCCTACggtcATtCAGC')

with open("out.extention.low.txt","w") as out:

for v in r.values():

out.write( "\t".join(v) +"\n")
